# β-Actin and Nuclear Myosin I are responsible for nucleolar reorganization during DNA Repair

**DOI:** 10.1101/646471

**Authors:** Elena Cerutti, Laurianne Daniel, Lise-Marie Donnio, Damien Neuillet, Charlene Magnani, Pierre-Olivier Mari, Giuseppina Giglia-Mari

## Abstract

During DNA Repair, ribosomal DNA and RNA polymerase I (rDNA/RNAP1) are reorganized within the nucleolus. Until now, the proteins and the molecular mechanism governing this reorganisation remained unknown.

Here we show that Nuclear Myosin I (NMI) and Nuclear Beta Actin (ACTβ) are essential for the proper reorganisation of the nucleolus, after completion of the DNA Repair reaction.

In NMI and ACTβ depleted cells, the rDNA/RNAP1 complex can be displaced at the periphery of the nucleolus after DNA damage but cannot re-enter within the nucleolus after completion of the DNA Repair. Both proteins act concertedly in this process. NMI binds the damaged rDNA at the periphery of the nucleolus, while ACTβ brings the rDNA back within the nucleolus after DNA repair completion. Our results reveal a previously unidentified function for NMI and ACTβ and disclose how these two proteins work in coordination to re-establish the proper rDNA position after DNA repair.

## Results and Discussion

Ribosome biogenesis is one of the most complex and energetically costly activities of the cell. The first and limiting step of ribosome biogenesis is the production of ribosomal RNA (rRNAs), specifically transcribed from ribosomal genes (rDNA) by the RNA polymerase I (RNAP1) within a specialized nuclear domain: the nucleolus. Despite nucleoli do not have membranes to isolate them from the nucleoplasm, they are impermeable to different nuclear proteins and their DNA content is kept inside during the majority of the cell cycle. Because of this apparent hermetical nature, some cellular functions, such as DNA repair of ribosomal genes imply that the DNA confined in the nucleolus would be externalised to allow repair proteins, and more generally, nuclear proteins, to access rDNA. Indeed, rDNA displacement at the periphery of the nucleolus happens during DNA replication (*1*) and DNA repair (*2–5*). Particularly, during DNA repair of UV lesion (*5*), rDNA and RNAP1 are displaced at the periphery of the nucleolus after UV-irradiation (displacement) and are repositioned within the nucleolus when DNA repair is completed (*5*) (repositioning) (Figure 1). Interestingly, these rDNA/RNAP1 movements are triggered by the presence of UV lesions on the DNA contained within the nucleolus and if UV damage is not fully repaired, the rDNA/RNAP1 complex remains at the periphery of the nucleolus (*5*).

**Figure 1.**
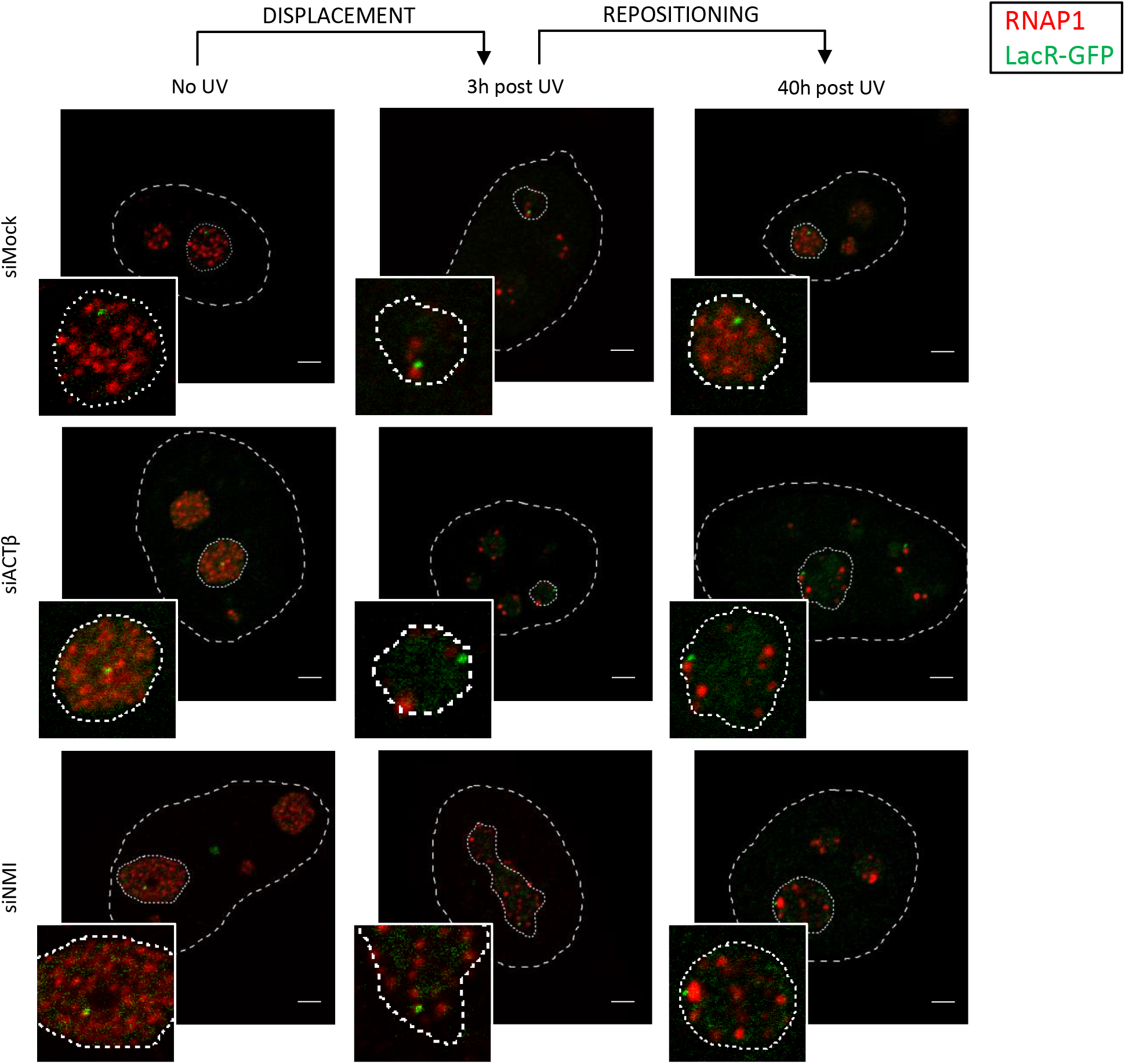
rDNA/RNAP1 repositioning is ACTβ and NMI-dependent. Confocal images of immunofluorescence assay against RNAP1 (red) in LacR-GFP (green) expressing cells transfected with siRNA against indicated factors and treated or not with UV-C. Nuclei and nucleoli are indicated by dashed lines and dotted lines respectively. Scale bar represents 3 µm.

The proteins involved in this displacement/repositioning cycle remain still unknown, as well as the molecular mechanism governing this movement. To disclose the foundation of this phenomenon, we explored the possibility that nuclear motors proteins could be responsible for the displacement and/or the repositioning of the rDNA/RNAP1 complex during DNA repair reactions. Two of the major nuclear motor proteins are nuclear β-actin (ACTβ) and Nuclear Myosin I (NMI). Firstly identified in the cytoplasm, ACTβ and NMI are also involved in several cellular events such as cell migration, muscle contraction or organelle movements (*5*) and interestingly these proteins have been implicated in long-range chromosomes movements within the nucleus (*6*), repositioning of active genes (*7*) and more recently in relocalisation of double strand breaks damaged DNA from the heterochromatin compartment to the nuclear periphery in Drosophila cells (*8*). Intriguingly, ACTβ and NMI are also involved in RNAP1 transcription (*9–11*), making them the best candidates to start exploring the rDNA/RNAP1 relocalisation during DNA repair.

We knocked down ACTβ and NMI in human fibroblasts (Figure S2) and UV irradiate them to induce the relocation of the rDNA/RNAP1 complex. RNAP1 was visualised by immunofluorescence staining while rDNA was visualised using a LacO-LacR-GFP system (kindly provided by W. Bickmore) in which Lac-O sequences were inserted in the close proximity of rDNA genes (Figure S1A). As previously observed, in siMock-treated cells rDNA and RNAP1 are displaced at the periphery of the nucleolus after DNA damage induction and are repositioned within the nucleolus when DNA repair is completed (*5*). In ACTβ and NMI depleted cells while the displacement of the rDNA/RNAP1 is unaltered, the repositioning is severely affected (Figure 1). Concomitantly, we measured the RNAP1 activity by measuring the 47S production by RNA-Fish (Figure S3, panel A and B). In our experimental conditions, depletion of ACTβ and NMI does not modify RNAP1 productivity, probably because depletion is not complete and the remaining 10-20 % of proteins (Figure S2) is sufficient to maintain an efficient RNAP1 transcription. Interestingly, after DNA repair completion, RNAP1 transcription restarts in ACTβ and NMI depleted cells as in siMock-treated cells. This situation was observed also in DNA-repair deficient cells (XP-C cells), in which UV-lesions on untranscribed DNA are not repaired, while transcribed sequences are repaired (*5*). In XP-C cells, as in ACTβ and NMI depleted cells, rDNA/RNAP1 remains at the periphery of the nucleolus because of remaining UV lesions on untranscribed sequences, while RNAP1 transcription restarts, as transcribed rDNA sequences are specifically repaired by the transcription coupled repair pathway (*5*). To exclude that depletion of ACTβ and NMI would induce a DNA repair deficiency, we conducted the Unscheduled DNA synthesis (UDS) measure, a specific assay to monitor the efficiency of the Global Genome Nucleotide Excision Repair pathway (GG-NER) which corrects UV-lesions on untranscribed DNA sequences and that is deficient in XP-C cells (Figure S5) (*5*). We could show that depleted ACTβ and NMI cells are proficient in GG-NER and that hence, UV-lesions are efficiently repaired, indicating that ACTβ and NMI are involved in the repositioning of the rDNA/RNAP1 after DNA repair completion. We wondered whether ACTβ and NMI involvement in this relocation was specific to the DNA repair reaction or if it was a general role in relocation of the rDNA/RNAP1 throughout other cellular stress events, such as transcription inhibition. To verify this, we treated ACTβ and NMI depleted cells with cordycepin to specifically induce RNAP1 transcription inhibition, the advantage of using cordycepin is that the effect is reversible simply by chasing it with a cordycepin-free medium, reproducing the displacement/repositioning cycle observed after UV-dependent DNA repair. We could show that ACTβ and NMI depleted cells are, in this case, efficient in both displacement and repositioning of rDNA/RNAP1 (Figure S7). These results show that ACTβ and NMI are specifically involved in the repositioning of rDNA/RNAP1 within the nucleolus after completion of the DNA repair reaction.

To investigate in details the implication of ACTβ and NMI in this process, we measured the binding activity of ACTβ and NMI on rDNA sequences by ChIP-qPCR using specific set of primers that locate along the rDNA genes and on the adjacent sequences outside of the transcribed rDNAs. We could perform this assay in absence of DNA damage, after UV irradiation during repair reaction and after completion of DNA repair (Figure 2A and 2B) and we could show that in absence of DNA damage, no detectable ACTβ and NMI are binding directly the rDNA genes or the adjacent untranscribed sequences. Remarkably, 3 hours after UV-irradiation, when RNAP1 transcription is shut down and the rDNA/RNAP1 is displaced at the periphery of the nucleolus both ACTβ and NMI bind rDNA genes and adjacent sequences, showing that the presence of DNA lesions and/or the position of rDNAs at the periphery of the nucleolus triggers the binding of ACTβ and NMI. Interestingly, when repair is completed (40 hours after UV irradiation), while NMI is released from rDNA (Figure 2A), ACTβ remains strongly bound to rDNA (Figure 2B). Intrigued, by the fact that NMI binds rDNAs at 3 hours post-irradiation but is released when DNA repair is completed and knowing that some research groups found NMI interacting with γH2AX chromatin (*12*), we explored the possibility that DNA damage signalling on rDNAs could be the trigger of the proper binding of NMI. In order to verify this hypothesis, we analysed rDNA/RNAP1 displacement and repositioning after UV-irradiation and DNA repair in cells treated with an ATR-inhibitor (Figure 3), which will impede UV-dependent phosphorylation of H2AX. Our results show that in absence of γH2AX phosphorylation the displacement of the rDNA/RNAP1 is unaltered while the repositioning is severely affected, exactly as in NMI depleted cells (Figure 3).

**Figure 2.**
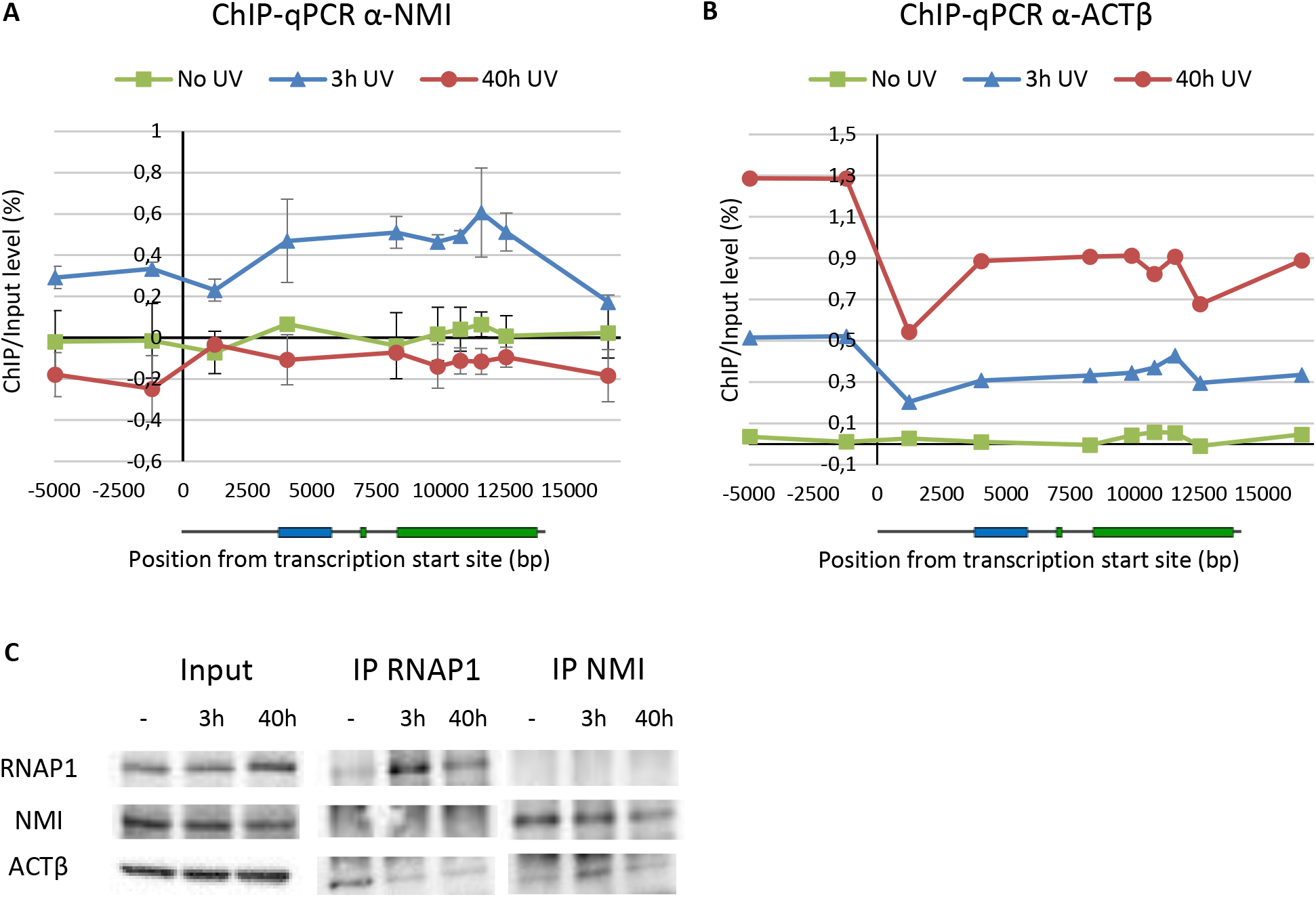
ACTβ and NMI binding activity on rDNA sequences and interaction with RNAP1. **A)** ChIP-qPCR results showing the binding profile of NMI on ribosomal DNA in MRC5 cells treated or not with UV-C. The y-axis depicts the ChIP/Input ratio minus background (Mock/Input ratio). Scqle bqr represents the SD **B)** ChIP-qPCR results showing the binding profile of ACTβ on ribosomal DNA in MRC5 cells treated or not with UV-C. The y-axis depicts the ChIP/Input ratio minus background (Mock/Input ratio). **C)** Chromatin Immunoprecipitation of RNAP1, NMI in MRC5 cells treated or not with UV-C. Bound proteins were revealed by Western Blot using antibodies against RNAP1, NMI and ACTβ. INPUT corresponds to 20% of the lysate used for IP reactions.

**Figure 3.**
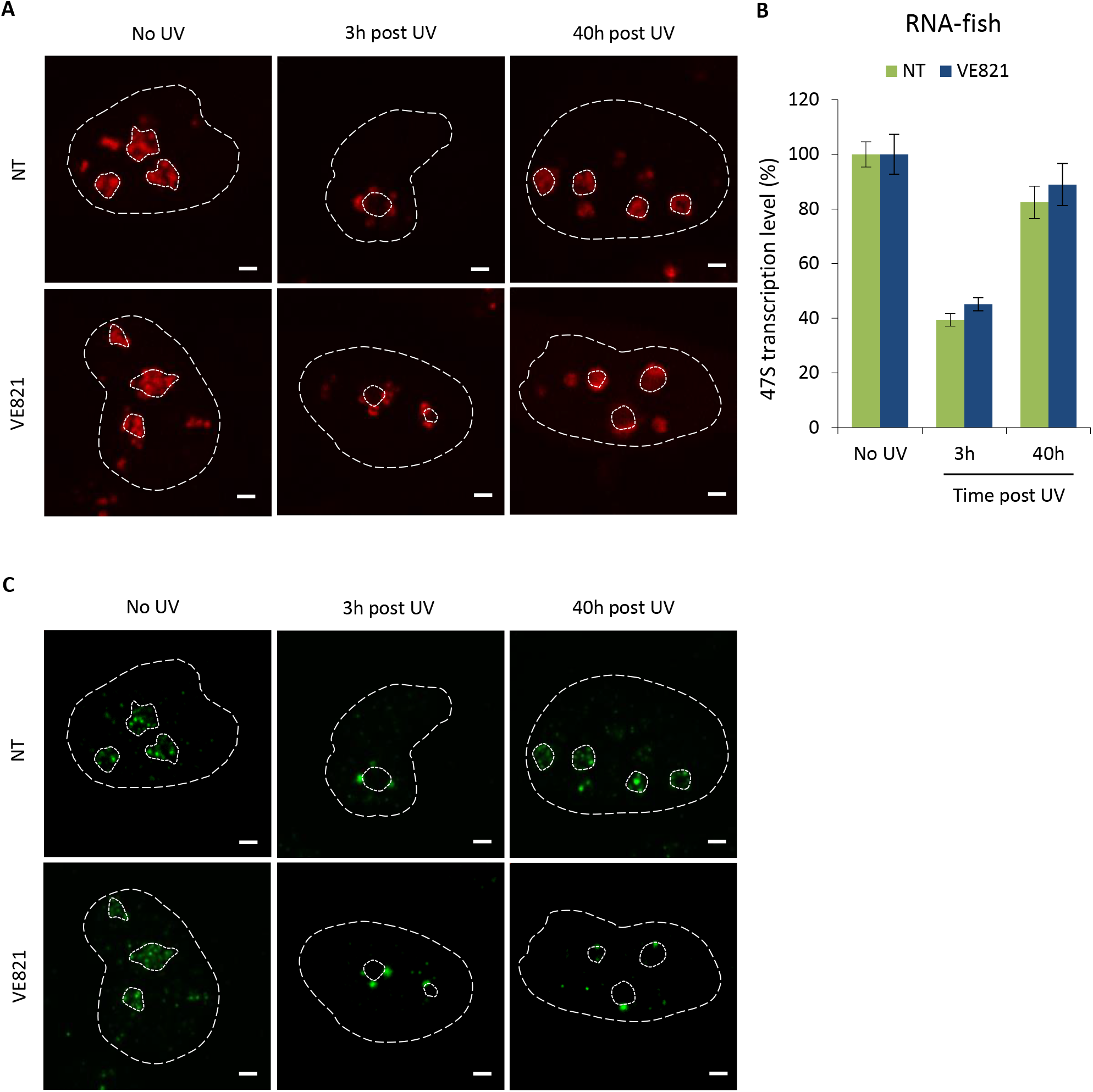
Impaired rDNA/RNAP1 post-UV repositioning upon ATR inhibition. **A)** Confocal images of RNA FISH labelling (red) performed on MRC5 cells treated or not with VE821 at 10 µM and treated or not with UV-C. Nuclei and nucleoli are indicated by dashed and dotted lines respectively. Scale bar represents 3 µm. **B)** Quantification of the RNA FISH experiments shown in A). Error bar represents the SEM. **C)** Confocal images of immunofluorescence assay against RNAP1 (green) in MRC5 cells treated or not with VE821 at 10µM and treated or not with UV-C. Nuclei and nucleoli are indicated by dashed lines and dotted lines respectively. Scale bar represents 3 µm.

To investigate whether ACTβ and NMI, interacts with RNAP1 or with each other during this process, we performed IP with RNAP1 and NMI antibodies. We could show that, in our experimental setting, RNAP1 interacts with ACTβ, more strongly in absence of DNA damage (Figure 2C) when RNAP1 transcription is not inhibited and rDNA/RNAP1 is located within the nucleolus. Interestingly, NMI interacts with ACTβ more strongly during DNA repair reactions when RNAP1 transcription is inhibited and the rDNA/RNAP1 is at the periphery of the nucleolus (Figure 2C). In our experimental setting, we could not find a direct interaction between NMI and RNAP1. These results suggest that ACTβ and NMI could work synergistically to bind rDNA and that both are needed in the same process of repositioning rDNA/RNAP1 within the nucleolus once repair is completed.

We propose a mechanistic model from the rDNA/RNAP1 repositioning (Figure 4) in which after UV exposure, rDNA/RNAP1 is displaced to the periphery of the nucleolus in order to allow repair proteins to access the lesion. The UV-induced phosphorylation of γH2Ax induces the binding of NMI to the rDNAs at 3 h post-irradiation and probably this interaction stimulates ACTβ molecules to be recruited on the damaged rDNAs sequences. After repair completion, while NMI is released from the rDNAs, more ACTβ molecules are bound on the repaired rDNAs sequences. In absence of ACTβ or NMI (or in absence of the γH2Ax signal) the rDNA/RNAP1 repositioning cannot take place.

**Figure 4.**
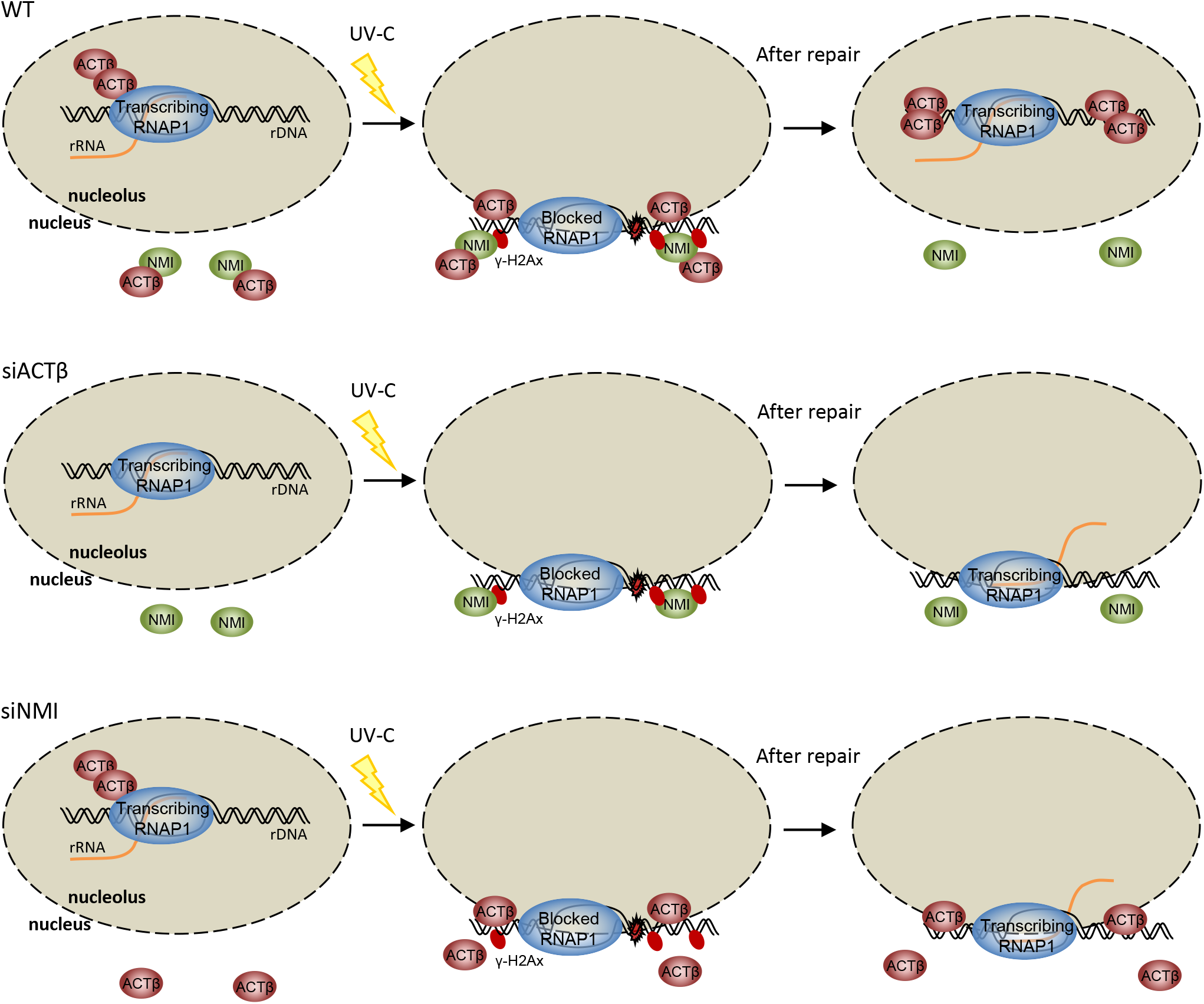
Model of the proposed mechanism involved in rDNA/RNAP1 repositioning. ACTβ binds RNAP1 during transcription of rDNA and NMI weakly binds ACTβ outside the nucleolus. After UV exposure, rDNA/RNAP1 is displaced to the periphery of the nucleolus in order to allow repair proteins to access the lesion. The UV-induced phosphorylation of γ-H2Ax induces the binding of NMI to the rDNAs at post-UV and probably this interaction stimulates ACTβ molecules to be recruited on the damaged rDNAs sequences. After repair completion, while NMI is released from the rDNAs, more ACTβ molecules are bound on the repaired rDNAs sequences. In absence of ACTβ or NMI (or in absence of the γ-H2Ax signal) the rDNA/RNAP1 repositioning cannot take place.

This work reveals ACTβ, NMI and the γH2Ax signal involvement in the rDNA/RNAP1 repositioning of rDNA/RNAP1 during repair of UV lesions. This is the starting point for further studies that will disclose the molecular mechanism of nucleolar motions. Many factors remain to be discovered, as well as the chromatin remodelling and the genomic environment of rDNA during and after this reorganization.

## Acknowledgements

We acknowledge Elizabeth Kerr and Jonathan Chubb (Wendy Bickmore’s research team, MRC Institute of Genetics & Molecular Medicine, Edinburgh, UK) for kindly providing the HT80 cells carrying the LacO/LacR system as well as all the information related to this cell line. This work was supported by La Ligue Nationale Contre le Cancer (LNCC), l’Agence Nationale de la Recherche (ANR FreTNET: ANR10-BLAN-1231-01; ANR DyReCT: ANR-14-CE10-0009) and the ARC foundation (Association pour la Recherche sur le Cancer).

## AUTHOR CONTRIBUTIONS

GGM and EC designed the experiments. EC, LD and LMD performed the experiments and analysed the data. POM assisted with the microscopy imaging and analysis. GGM, EC and LD wrote the paper.

## Conflicts of interest

The authors disclose no potential conflict of interest.

## Extended Data

**Figure S1.**
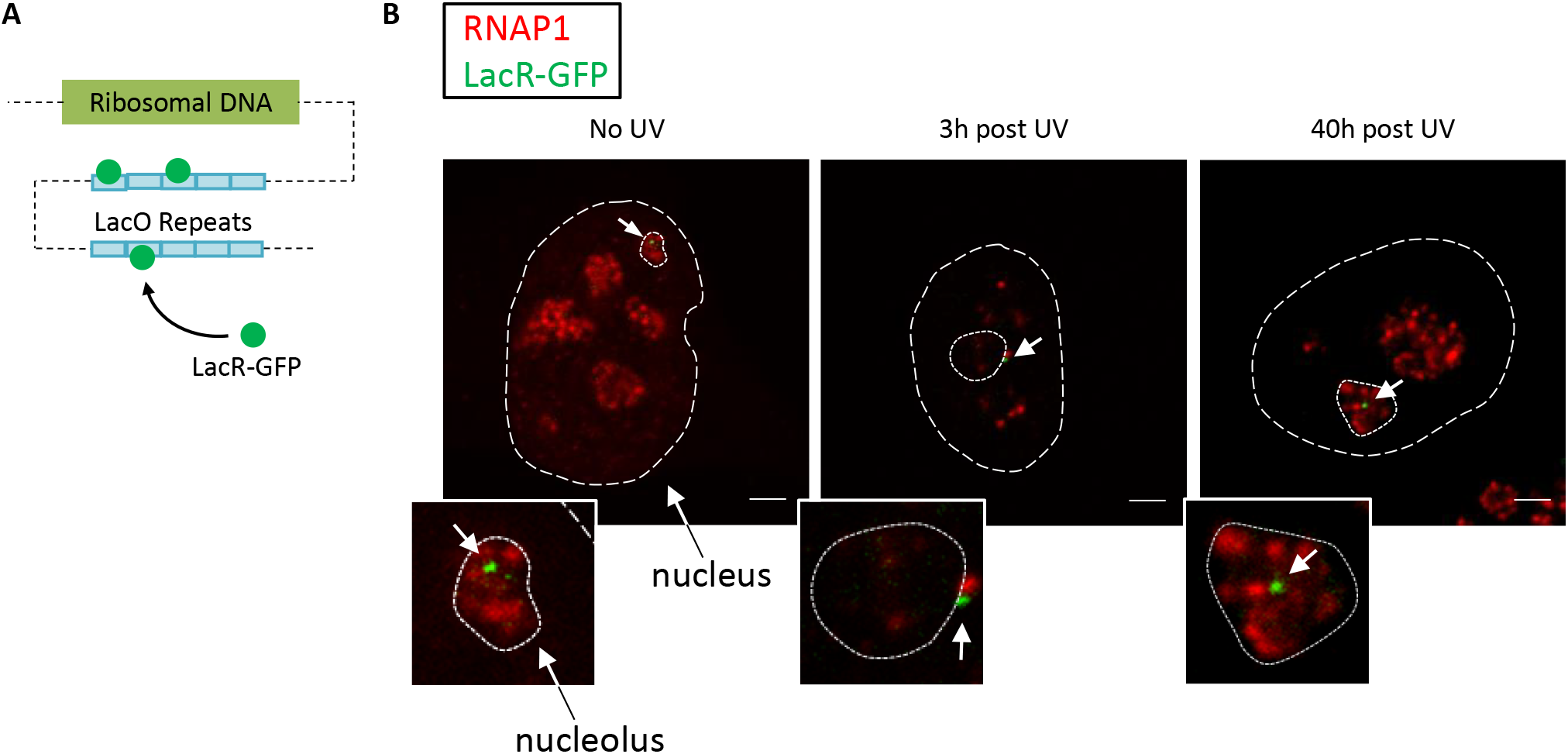
rDNA location during DNA repair. **A)** Schematic representation of the LacO/LacR system used to visualize the ribosomal DNA. Several Lac Operon genes were integrated downstream of the rDNA of the cells, which were also transfected with a plasmid carrying the Lac Repressor gene tagged with the GFP coding sequence. **B)** Confocal images of immunofluorescence assay against RNAP1 (red) in LacR-GFP (green) expressing cells treated or not with UV-C. Nuclei, nucleoli and LacO array (GFP green dot) are indicated by dashed lines, dotted lines and white arrows respectively. Scale bar represents 3 µm.

**Figure S2.**
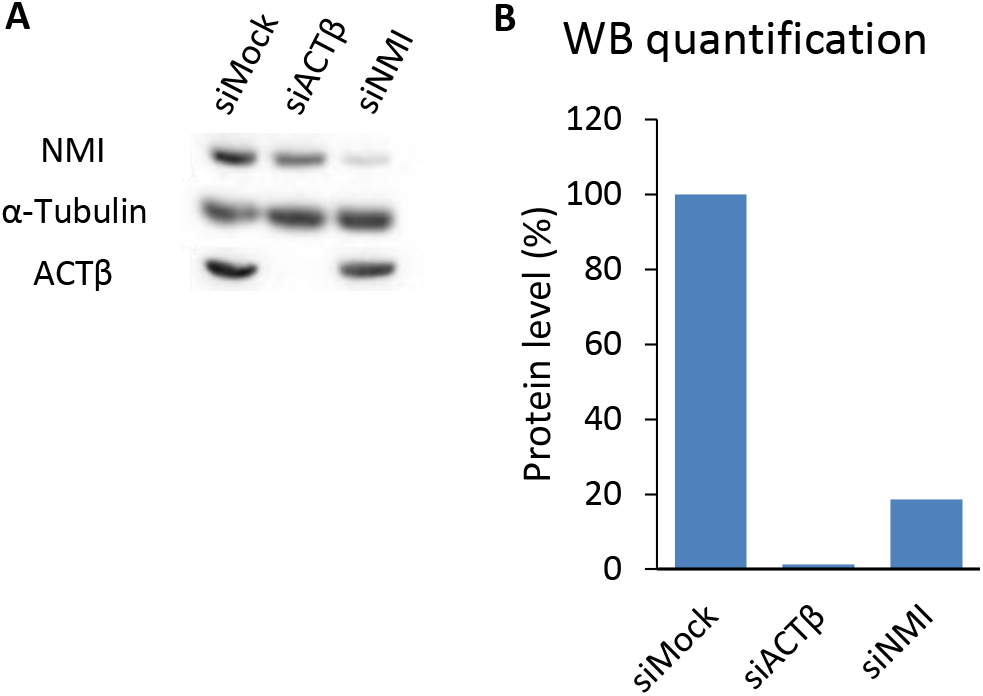
**A)** Western Blot on whole cell extracts performed on LacO/LacR-GFP cells treated with siRNA against indicated factors and represented in Figure 1. **B)** Quantification of WB shown in A). Protein amount is normalized to α-Tubulin signal.

**Figure S3.**
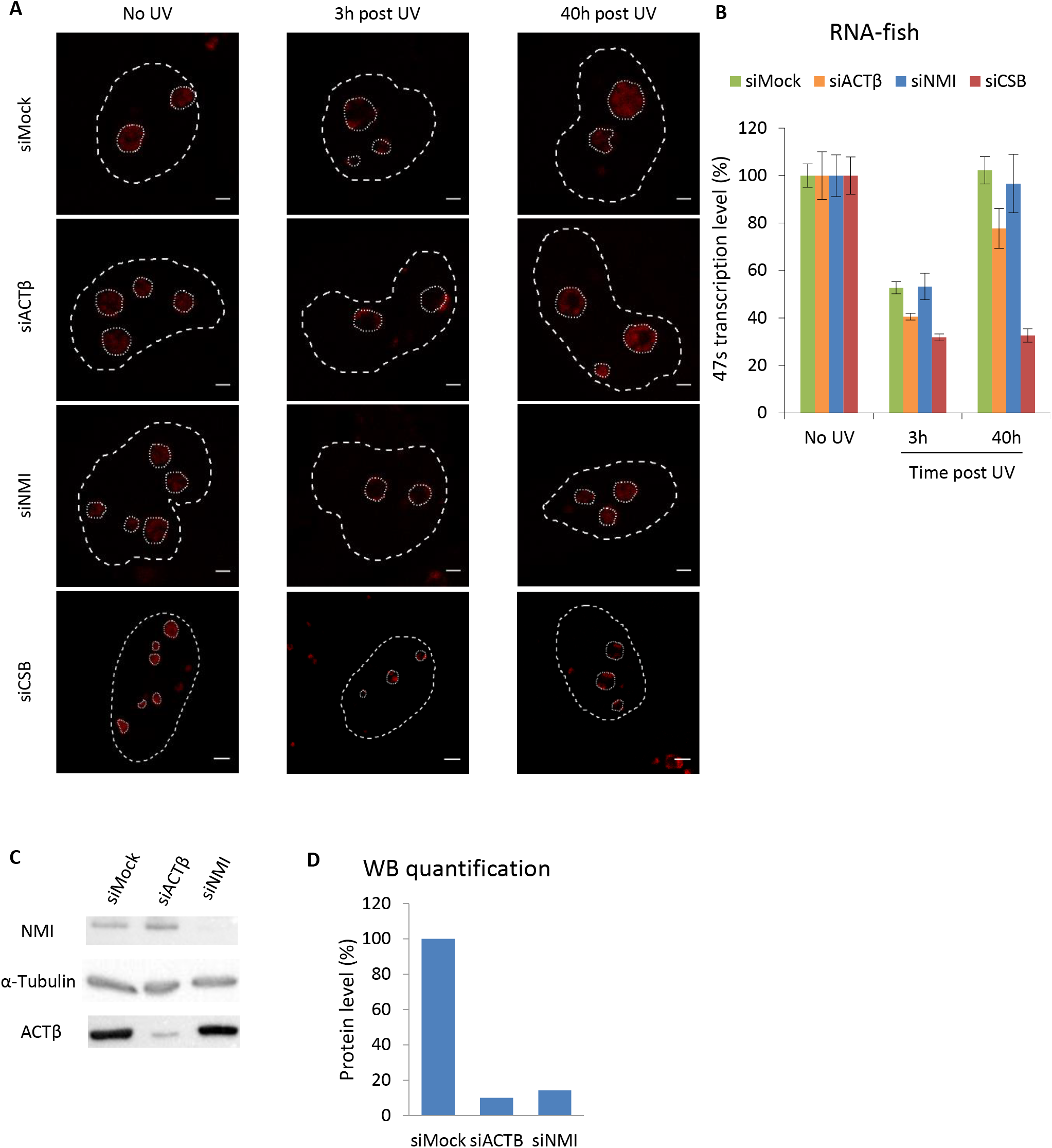
**A)** Confocal images of RNA FISH labelling (red) performed on MRC5 cells transfected with siRNA against indicated factors and treated or not with UV-C. Nuclei and nucleoli are indicated by dashed and dotted lines respectively. Scale bar represents 3 µm. **B)** Quantification of the RNA FISH experiments shown in A). Error bar represents the SEM. **C)** Western Blot on whole cell extracts performed on MRC5 cells of RNA FISH experiment shown in A) **D)** Quantification of WB shown in C). Protein amount is normalized to α-Tubulin signal.

**Figure S4.**
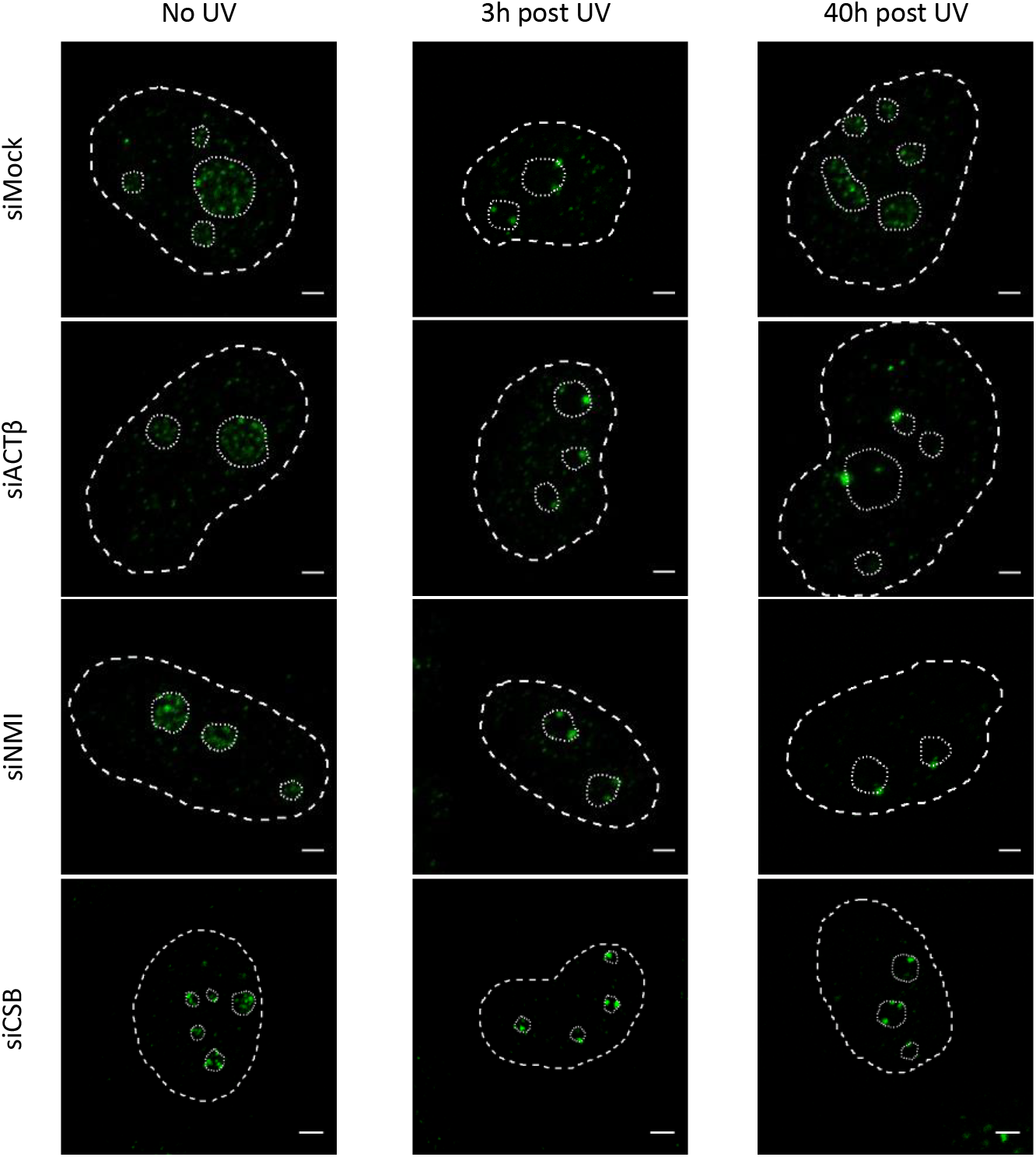
Confocal images of immunofluorescence assay against RNAP1 (green) in MRC5 cells transfected with siRNA against indicated factors and treated or not with UV-C. Nuclei and nucleoli are indicated by dashed lines and dotted lines respectively. Scale bar represents 3 µm.

**Figure S5.**
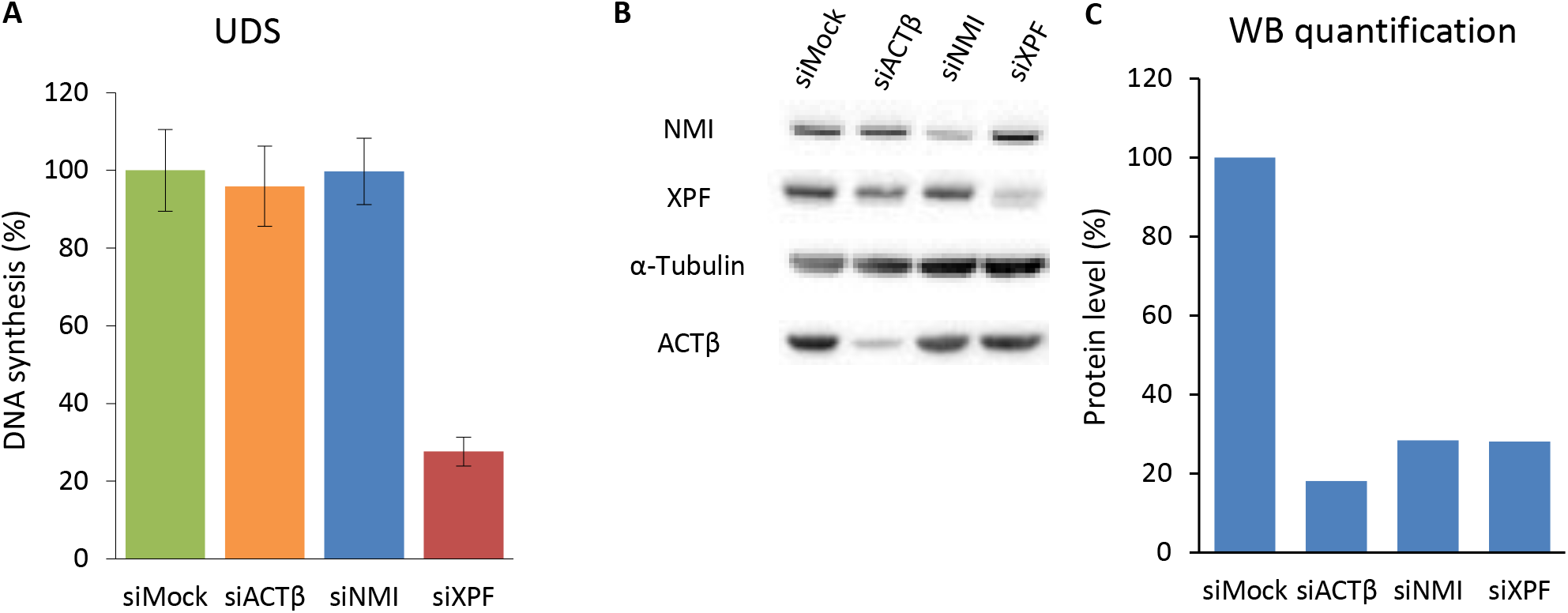
**A)** Unscheduled DNA Synthesis determined by EdU incorporation after local damage induction with UV-C in MRC5 cells treated with siRNAs against indicated factors. **B)** Western Blot on whole cell extracts performed on MRC5 cells of UDS experiment shown in A). **C)** Quantification of WB shown in B). Protein amount is normalized to α-Tubulin signal.

**Figure S6.**
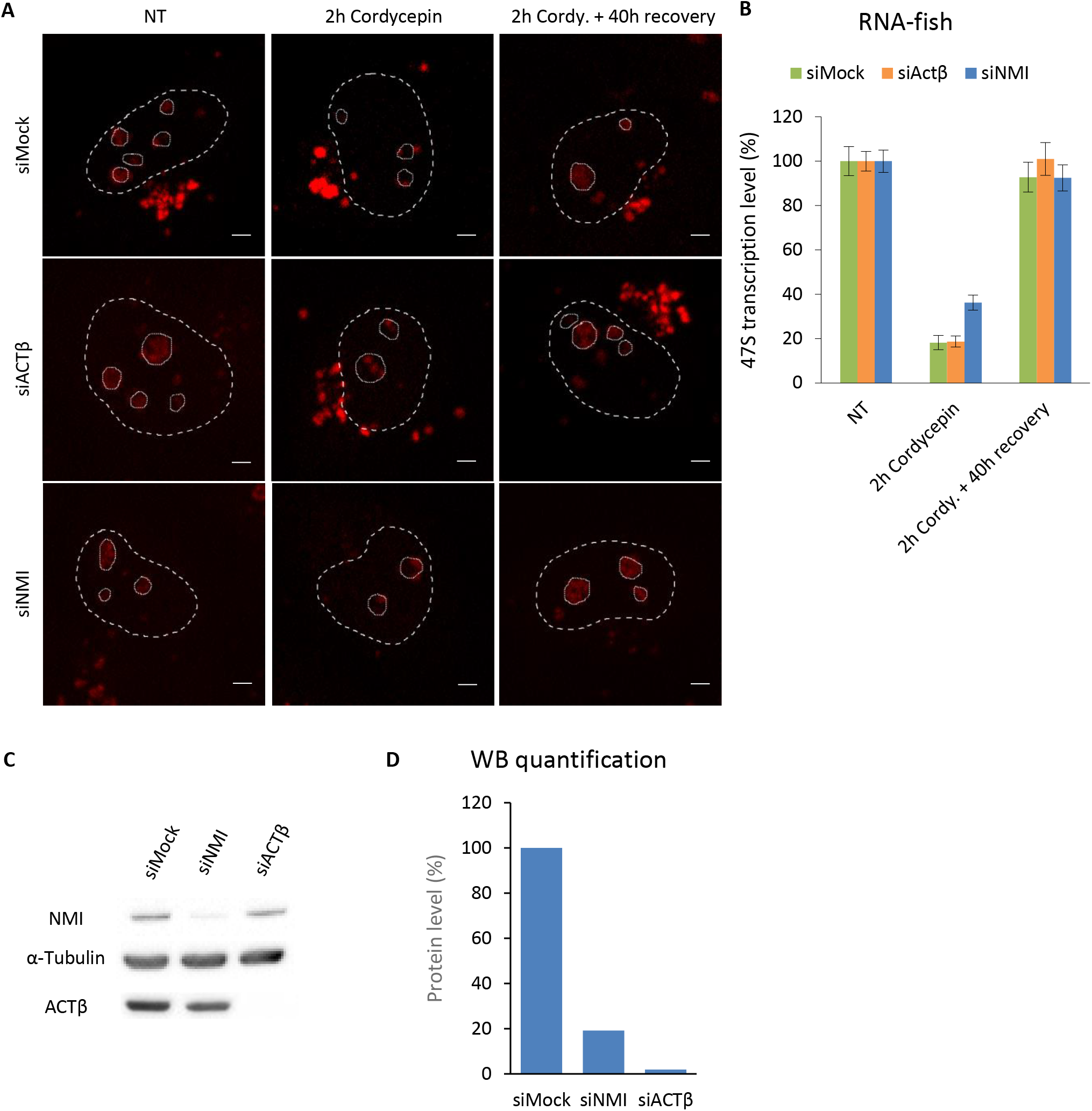
**A)** Confocal images of RNA FISH labelling (red) performed on MRC5 cells transfected with siRNA against indicated factors and treated or not with Cordycepin at 50 µg/ml. Nuclei and nucleoli are indicated by dashed and dotted lines respectively. Scale bar represents 3 µm. **B)** Quantification of the RNA FISH experiments shown in A). Error bar represents the SEM. **C)** Western Blot on whole cell extracts performed on MRC5 cells of RNA FISH experiment shown in A) **D)** Quantification of WB shown in C). Protein amount is normalized to α-Tubulin signal.

**Figure S7.**
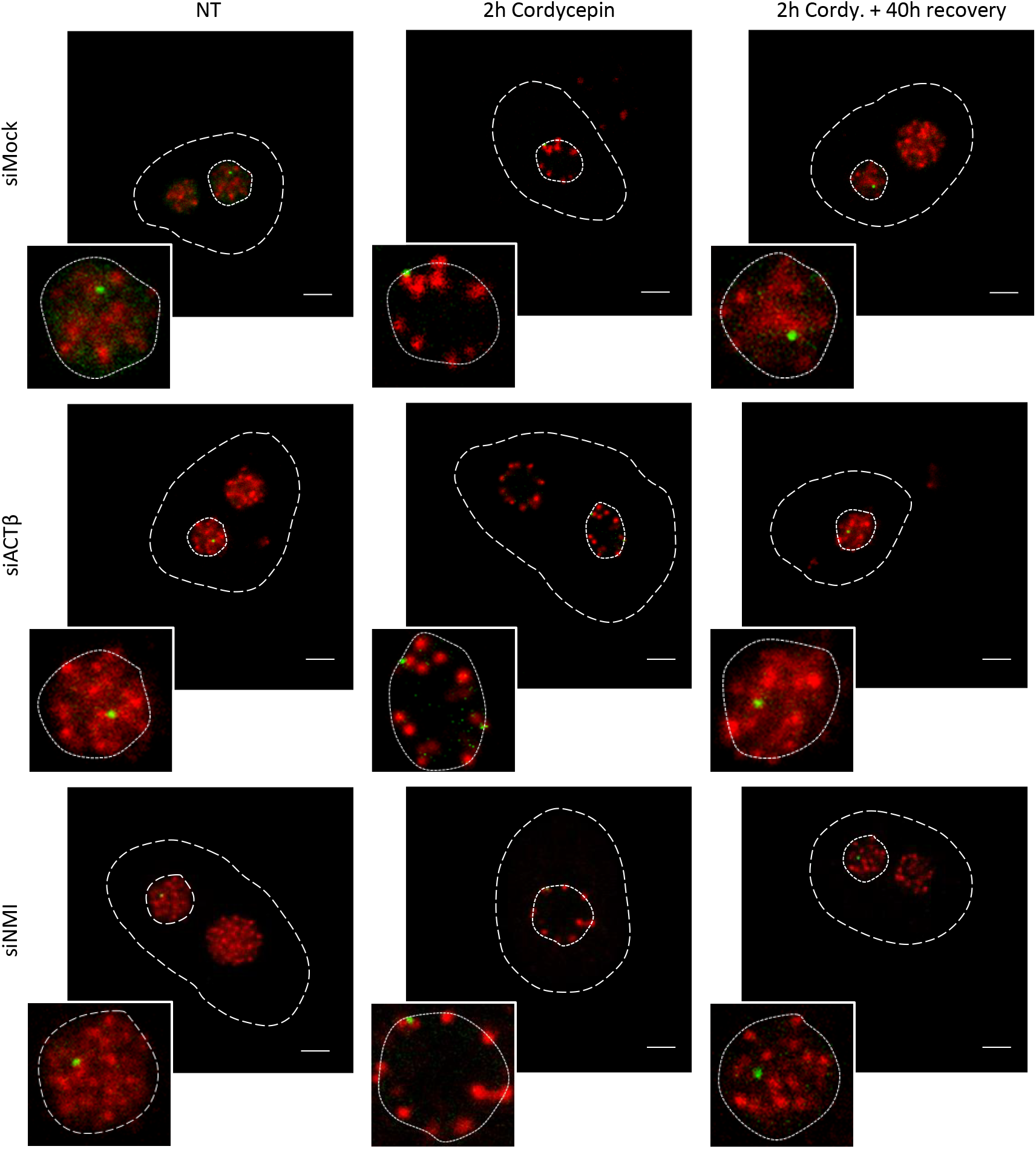
Confocal images of immunofluorescence assay against RNAP1 (red) in LacR-GFP (green) expressing cells transfected with siRNA against indicated factors and treated or not with Cordycepin at 50 µg/ml. Nuclei and nucleoli are indicated by dashed lines and dotted lines respectively. Scale bar represents 3 µm.

## Methods

### Cell culture and treatments

Wild type SV40-immortalized human fibroblasts (MRC5) were cultured in a 1:1 mixture of Ham’s F10 and DMEM (Lonza) supplemented with 1% antibiotics (penicillin and streptomycin; Lonza) and 10% fetal bovine serum (Gibco) and incubated at 37°C with 20% O_2_ and 5% CO_2_.

HT-1080 cells (Rasheed et al., 1974) stably expressing an adapted Lac Operator/Lac Repressor (LacO/LacR) system (selected using BlasticidinS and Hygromycin, 5 µg/ml and 100 µg/ml respectively), were used to detect the rDNA as previously described (Chubb et al., 2002; Robinett et al., 1996). These cells were cultured in DMEM (Lonza), supplemented with 1% antibiotics (penicillin and streptomycin; Lonza) and 10% fetal bovine serum (Gibco), and at 37°C with 20% O_2_ and 5% CO_2_.

DNA damage was inflicted by UV-C light (254 nm, 6-Watt lamp). Cells were globally irradiated with a 16 J/m^2^ dose of UV-C or locally irradiated with a 100 J/m^2^ dose of UV-C through a Millipore filter (holes of 5 µm of diameter). Experiments were performed at different time points after UV exposure (3h and 40h post UV). Not irradiated cells (No UV) were used as control.

RNAP1 transcription inhibition has been achieved by 2h incubation in medium containing Cordycepin at 50 µg/ml. Resumption of transcription has been obtained by replacement of Cordycepin medium with normal medium for 40h. Not treated cells (NT) were used as control.

VE821 drug was used at the concentration of 10 µM. Cells were treated in VE821 containing medium for 3h, then cells were UV-C globally irradiated with a 16 J/m^2^ dose, or not irradiated as control, and left in drug containing medium for 40h or 3h before fixation.

### Transfection of small interfering RNAs (siRNAs)

On day 0, 100 000 cells were seeded in a 6-wells plate and/or on 18 mm coverslips. The first and second transfections were performed on day 1 and day 2, using Lipofectamine® RNAiMAX reagent (Invitrogen; 13778150) or Gen Jet (Tebu-Bio), according to the manufactures’ protocols. Experiments were performed between 24h and 72h after the second transfection. SiRNA efficiency was confirmed by western blot on whole cell extracts. SiRNAs sequences are described in Table 1.

**Table 1.**
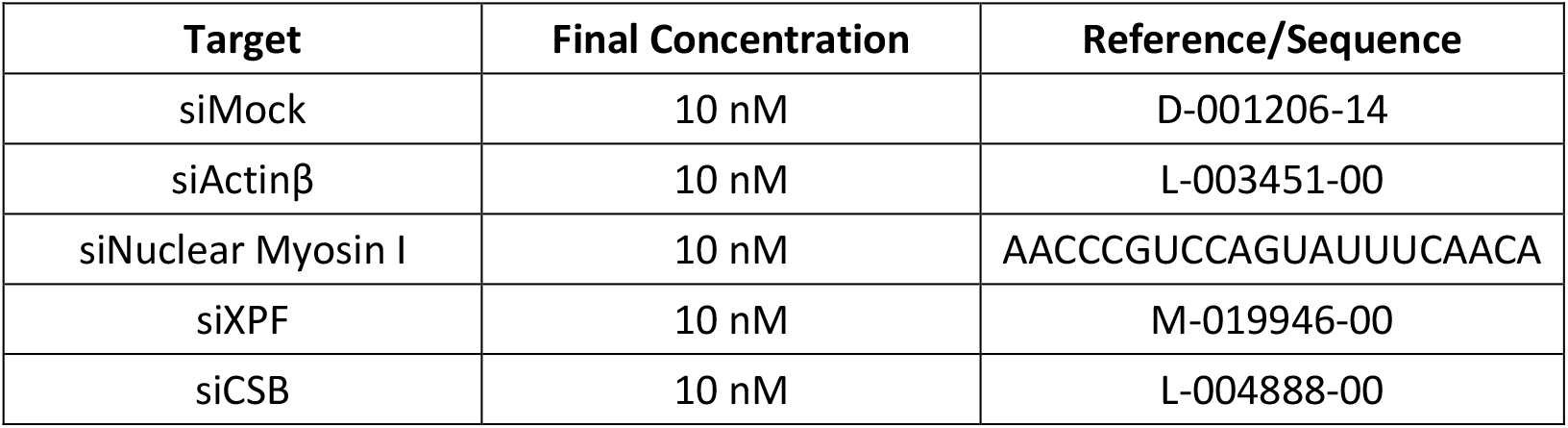
Small interfering RNAs

### Whole cell extracts

Cells were collected using trypsin and centrifuged 10 min at 1400 rpm. Firstly, cell pellet was washed with PBS supplemented with the Protease Inhibitor Cocktail (Roche) and spinned down 10 min at 1400 rpm. Secondly, cell pellet was incubated with Lysis buffer (ProteoJETTM Mammalian Cell Lysis Reagent, Fermentas) complemented with the Protease Inhibitor Cocktail (Roche), for 10 min at room temperature on a shaker (500 rpm). Finally, samples were centrifuged at 16000 g for 15 min and supernatant was freezed at - 80° C. Protein concentration was determined using the Bradford method, samples were diluted with 1X Laemmli buffer and heated 10 min at 95° C.

### Chromatin extracts

All procedures were carried out on ice unless otherwise stated. Cells were grown in 14.5 cm dish. After treatments, cells were washed twice with PBS and cross-linked with a solution of 1 % formaldehyde in PBS (7.5 min at RT, shaking) prepared from a 37 % stock (Sigma-Aldrich, F1635). Cross-linking was neutralized by adding glycine for a final concentration of 0.125 M, followed by a wash with cold PBS. Cells were collected by scraping in PBS supplemented with the EDTA-free protease inhibitor tablets (Roche) and centrifuged 10 min at 2000 rpm and 4° C.

All buffers used for chromatin extraction contained, among others, the EDTA-free protease inhibitor tablets (Roche).

Cell pellet was suspended in Lysis buffer (50 mM Hepes-KOH [pH 8], 1 mM EDTA, 500 mM NaCl, 10 % glycerol, 0.5 % NP-40, 0.25 % Triton X-100) and incubated 10 min rotating at 4° C. The suspension was centrifuged 10 min at 2000 rpm and 4° C. Cell pellet was then washed with Wash buffer (10 mM Tris-HCl [pH 8], 1 mM EDTA, 500 mM NaCl), incubated 10 min rotating at 4° C and centrifuged 10 min at 2000 rpm and 4° C. Cell pellet was finally incubated for 30 min in 1 ml IP buffer (50 mM Hepes-KOH [pH 7.5], 1 mM EDTA, 500 mM NaCl, 1 % Triton X-100, 0.1 % Na-deoxycholate, 0.1 % SDS) before sonication. The nuclear suspension was sonicated using the S220 Focused-ultrasonicator (Covaris) for 8 min (Average Incident Power 14 Watt, Peak Power 140 Watt, Duty Factor 10 %, Cycle/Burst 200 count) to yield DNA fragments with an average size of 250 bp. Samples were then centrifuged (14.000 rpm, 10 min, 4° C) and the supernatant containing the cross-linked chromatin was freezed at - 80° C. DNA concentration was quantified using the NanoDropTM 2000 Spectrophotometer (Termo Scientific).

### ChIP qPCR

50 µg of MRC5 chromatin extracts were incubated overnight at 4° C in 150 μl total volume of IP buffer (see above) with antibody (ACTβ: ab8227 Abcam; NMI: M3567 Sigma) (ChIP) or no antibody (Mock). Immunoprecipitation (IP) was performed for 1 hour with 40 μl of washed magnetic Bio-Adembeads Protein G (Ademtech). After IP, the chromatin-beads interaction was washed twice with IP buffer, once with Na-deoxycholate buffer (10 mM Tris-HCl [pH 8], 1 mM EDTA, 250 mM LiCl, 0.5 % NP-40, 0.5 % Na-deoxycholate) and once with TE1 buffer (50 mM Tris-HCl [pH 8], 10 mM EDTA). Chromatin was then eluted in Elution buffer (50 mM Tris-HCl [pH 8], 10 mM EDTA, 1 % SDS) for 20 min at 37° C, 1400 rpm and diluted in TE2 buffer (10 mM Tris-HCl [pH 8], 1 mM EDTA). DNA from ChIP, Mock and Input preparations were decrosslinked and purified by phenol chloroform extraction. Samples were amplified by real-time PCR (qPCR) using the Power SYBR Green PCR master mix (Applied Biosystems) on a CFX Connect™ Real-Time PCR Detection System (BioRad). ChIP data were normalized to the Input (to consider copy number) and subtracted with the background (Mock). Primer sequences for qPCR are listed in Table 2.

**Table 2.**
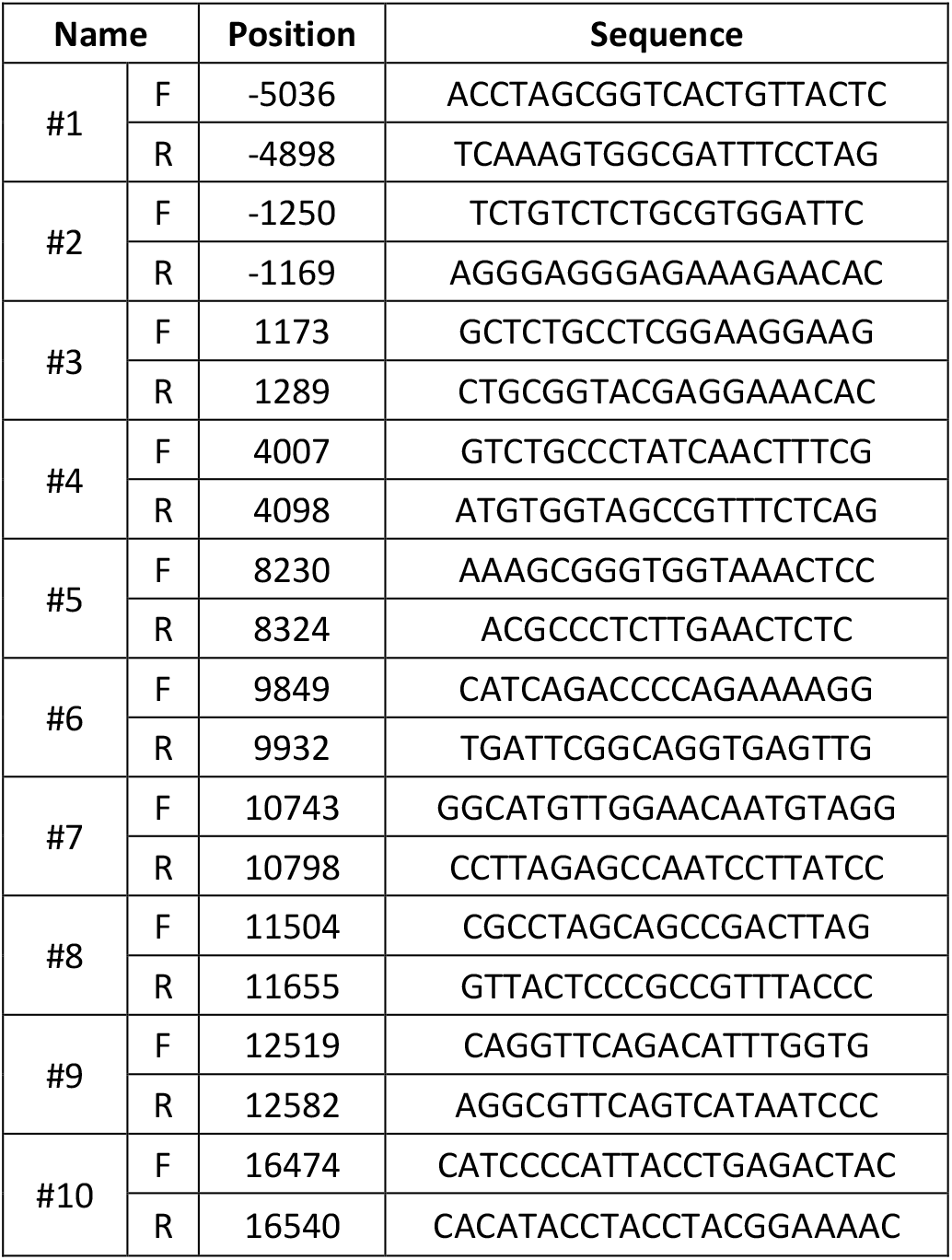
Primers used for ChIP-qPCR experiment

### ChIP WB

90 µg of MRC5 chromatin extracts were incubated overnight at 4° C in 150 μl total volume of IP buffer (see above) with antibody (RNAP1: anti-RPA194, sc-48385, Santa Cruz; NMI: M3567 Sigma) (ChIP) or no antibody (Mock). Immunoprecipitation was performed as for ChIP qPCR experiment (see above). Chromatin was then eluted with 2X Laemmli buffer. ChIP, Mock and Input preparations were heated at 95° C for 45 min and loaded on a SDS-PAGE gel.

### Western blot

Proteins were separated on a SDS-PAGE gel at an acrylamide percentage appropriate to protein size and transferred onto a polyvinylidene difluoride membrane (PVDF, 0.45 μm Millipore). The membrane was blocked in 5 % milk in PBS 0.1 % Tween 20 (PBS-T) solution and incubated for 1.5 h RT or overnight at 4° C with the primary antibodies in milk PBS-T (RNAP1: anti-RPA194, sc-48385, Santa Cruz; ACTβ: A5316, Sigma or ab8227, Abcam; NMI: M3567, Sigma; α-Tubulin: T5168, Sigma, XPF: MS-1351-P1, NeoMarkers). Subsequently, membrane was washed repeatedly with PBS-T and incubated 1 h RT with the secondary antibody in milk PBS-T (Goat anti-mouse IgG HRP conjugate (170-6516; BioRad); Goat anti-rabbit IgG HRP conjugate (170-6515; BioRad)). After the same washing procedure, protein bands were visualized via chemiluminescence (ECL Enhanced Chemo Luminescence; Pierce ECL Western Blotting Substrate) using the ChemiDoc MP system (BioRad).

### RNA Fluorescence In Situ Hybridization

Cells were grown on 18 mm coverslips, washed with warm (37°C) PBS and fixed with 4% paraformaldehyde for 15 min at 37° C. Coverslips were washed twice with PBS. Cells were permeabilized by washing with PBS 0.4 % Triton X-100 for 7 min at 4° C. Cells were washed rapidly with PBS before incubating them with pre-hybridization buffer (2X SSPE and 15 % formamide) (20X SSPE, [pH 8.0]: 3 M NaCl, 157 mM NaH2PO4.H2O and 25 mM EDTA) for at least 30 min. 3.5 µl of probe (10 ng/ml) was diluted in 70 µl of hybridization mix (2X SSPE, 15 % formamide, 10 % dextran sulphate, 0.5 mg/ml tRNA) and heated at 90° C for 1 min. Hybridization of the probe was conducted overnight at 37° C in a humidified environment. Subsequently, cells were washed twice for 20 min with pre-hybridization buffer, then once for 20 min with 1X SSPE and finally mounted with Vectashield (Vector Laboratories) and kept at −20° C. The probe sequence (5’ to 3’) is: Cy5-AGACGAGAACGCCTGACACGCACGGCAC. At least 30 cells were imaged for each condition.

### Immunofluorescence

Cells were grown on 18 mm coverslips, washed with warm (37° C) PBS and fixed with 2% paraformaldehyde for 15 min at 37° C. Cells were permeabilized with PBS 0.1 % Triton X-100 (3X short + 2X 10 min washes). Blocking of non-specific signal was performed with PBS+ (PBS, 0.5 % BSA, 0.15 % glycine) for at least 30 min. Then, coverslips were incubated with 70 µl of primary antibody mix (RNAP1: Mouse anti-RPA194, 1/500 in PBS+, sc-48385) for 2 h at RT in a moist chamber, washed with PBS (3X short + 2X 10 min), quickly washed with PBS+ before incubating with 70 µl of secondary antibody mix (Goat anti-mouse Alexa Fluor® 488 or Goat anti-mouse Alexa Fluor® 594 1/400 in PBS+, Invitrogen) for 1 h at RT in a moist chamber. After the same washing procedure, coverslips were finally mounted using Vectashield with DAPI (Vector Laboratories) and kept at - 20° C. At least 15 cells were imaged for each condition.

### Unscheduled DNA Synthesis (UDS)

MRC5-SV40 immortalized human fibroblasts, were grown on 18 mm coverslips. After local irradiation (100 J/m^2^ UV-C) through a 5 µm pore polycarbonate membrane filter, cells were incubated for 3 hours with ethynyldeoxyuridine, washed, fixed and permeabilized. Fixed cells were treated with a PBS-blocking solution (PBS+: PBS containing 0.15% glycine and 0.5% bovine serum albumin) for 30 min, subsequently incubated with primary antibodies mouse monoclonal anti-yH2AX (Ser139) (Upstate, clone JBW301) 1/500 diluted in PBS+ for 1h, followed by extensive washes with Tween20 in PBS. Cells were then incubated for 1h with secondary antibodies conjugated with Alexa Fluor 594 fluorescent dyes (Molecular Probes, 1:400 dilution in PBS+). Then, cells were incubated for 30 min with the Click-iT reaction cocktail containing Alexa Fluor Azide 488. After washing, the coverslips were mounted with Vectashield containing DAPI (Vector). Images of the cells were obtained with the same microscopy system and constant acquisition parameters. Images were analysed using ImageJ as follows: (i) a ROI outlining the locally damaged area was defined by using the yH2AX staining, (ii) a second ROI of comparable size was defined in the nucleus (avoiding nucleoli and other non-specific signals) to estimate background signal, (iii) the ‘local damage’ ROI was then used to measure the average fluorescence correlated to the EdU incorporation, which is an estimate of DNA synthesis after repair once the nuclear background signal obtained during step (ii) is subtracted. For each sample three independent experiments were performed.

### Fluorescent imaging and analysis

Imaging has been performed on a Zeiss LSM 780 NLO confocal laser-scanning microscope (Zeiss), using a 60x/1.4 oil objective. Images were analyzed with ImageJ software. For all images of this study, nuclei and nucleoli were delimited with dashed and dotted line respectively, using DAPI staining or transmitted light.

### Statistical analysis

Error bars represent the Standard Error of the Mean (SEM) of the biological replicates.

